# Single cell gene set scoring with nearest neighbor graph smoothed data (gssnng)

**DOI:** 10.1101/2022.11.29.518384

**Authors:** David L Gibbs, Michael K Strasser, Sui Huang

## Abstract

Gene set scoring (or enrichment) is a common dimension reduction task in bioinformatics that can be focused on differences between groups or at the single sample level. Gene sets can represent biological functions, molecular pathways, cell identities, and more. Gene set scores are context dependent values that are useful for interpreting biological changes following experiments or perturbations. Single sample scoring produces a set of scores, one for each member of a group, which can be analyzed with statistical models that can include additional clinically important factors such as gender or age. However, the sparsity and technical noise of single cell expression measures create difficulties for these methods, which were originally designed for bulk expression profiling (microarrays, RNAseq). This can be greatly remedied by first applying a smoothing transformation that shares gene measure information within transcriptomic neighborhoods. In this work, we use the nearest neighbor graph of cells for matrix smoothing to produce high quality gene set scores on a per-cell, per-group, level which is useful for visualization and statistical analysis.

**Availability and implementation:** The gssnng software is available using the python package index (PyPI) and works with Scanpy AnnData objects. It can be installed using ‘pip install gssnng’. More information and demo notebooks: See https://github.com/IlyaLab/gssnng

## Introduction

In biological systems, groups of genes carry out biological functions via pathways, protein complexes, and signalling cascades. It is often informative to assess the activity of these transcriptional programs through examining the concerted expression of several genes together. Their individual expression may be weak, but co-expression of genes in a pathway is often a strong indication of activity. Using gene set enrichment techniques can shed light on how pathways and modules take part in the response to perturbations [1, 2, 3, 4].

Recently, single sample methods have been developed to compute a gene set score independently for each sample. The matrix of scores (samples by gene set) are used as the basis of further analysis, such as in visualizations and statistical modeling [5, 6]. With single sample scores, statistical models can be better specified since they can include not only gene set scores but also clinical, biospecimen, and technical variables. From single sample analysis, it is a natural extension to single cells.

Much of the previously mentioned methods were designed for bulk expression data like gene microarrays and RNA-seq, where millions of cells are batch processed. These experiments provide ample measures on practically all genes, but represent an unknown mixture of cells (i.e. a population average). Single cell transcriptomics provides more precise information on the mixture of cell types, the heterogeneity of those cells, and allows for the discovery of new subtypes. However, the data is shallow (few counts), sparse (many zeros in the expression matrix), and noisy with a much smaller collection of total genes measured in each cell [7]. This causes difficulties in standard types of analysis, such as differential expression, but also gene set analysis. For example, methods that rely on ranked expression profiles, like ssGSEA, will be operating on data that have shared rank across the high proportion of zeros or integer collisions (e.g. genes with 1 count). Due to frequent ties, ranking becomes highly unstable; small changes in a gene’s counts lead to large changes in rank, further leading to large changes in gene set scores.

In the related task of determining differential expression, recent studies have shown that these problems can be avoided by using “pseudobulk” profiles, created by summing across groups of cells [8]. The summation is a dimensional reduction (in the cell dimension) that creates higher abundances, lower noise floors, breaks many of the ties in expression counts, and allows for better overlap with gene sets. However, in applying this ‘pseudo-bulk’ transformation, most of what makes single cell data valued is lost; namely the heterogeneity and variability observed across cells and conditions. Other approaches have been developed that address the noise and sparsity and in many instances these focus on K-nearest neighbor graphs, which are a fundamental part of single cell data analysis and has thus been an attractive target for data processing, such as in MetaCell [9] which partitions subgraphs to model an archetypal cell. Also, other methods using nearest neighbor smoothing have been reported to be highly effective in single cell data sets [10, 11]. Methods such as MAGIC [12] involve modeling the lower dimensional data manifold with diffusion on the KNN graph or in scGSEA [13] estimating latent spaces is used for gene set scoring.

With these approaches in mind, we have developed a python package that works with Scanpy AnnData objects to produce a gene set score for each cell [14]. Included are a collection of scoring functions from previously described single sample methods, similar to decoupleR [15], and functions that ingest gene sets from standard .gmt files or from OmniPath [16].

### 1. Overview of the method

Briefly, cells are grouped using user-specified attributes such as cell type, cluster label, batch, or condition. These groups are used to define collections of cells that form disjoint nearest neighbor graphs. After creating groups, the remainder of the process is done in parallel by group. The graphs, represented as a matrix, are used in smoothing the gene expression counts matrix. Lastly, the smoothed expression profile for each cell is passed to a selected gene set scoring function (Figure 1A).

**Fig. 1.**
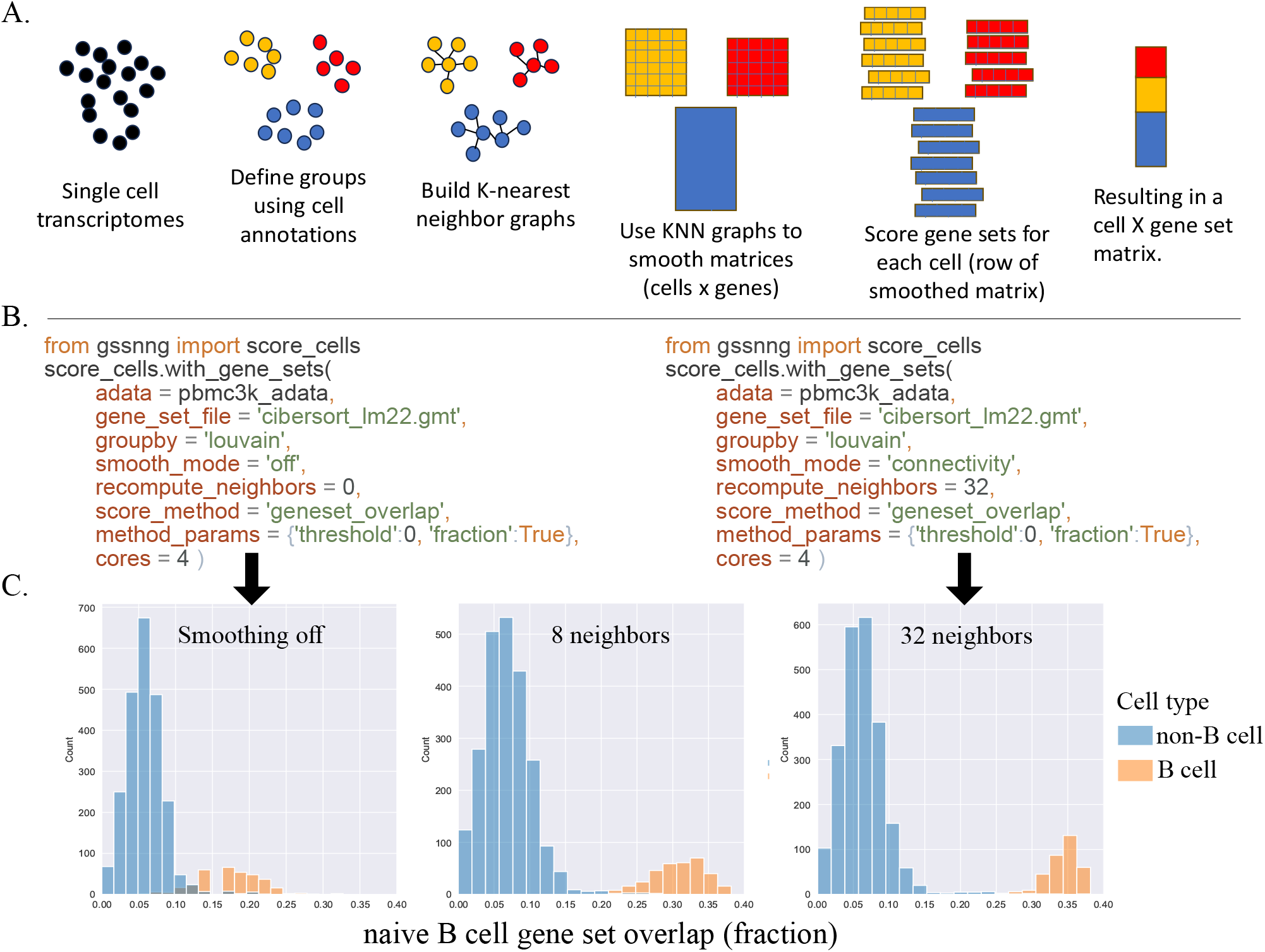
Figure 1. (A.) Overview of the approach. (B.) Code examples to generate the results shown in next panel. (C.) Effect of neighborhood smoothing for PBMCs and a B-cell signature. Gene set overlap (cells from 10X 3kPBMC dataset) with smoothing turned off, or K-nearest neighbor smoothing(K=8 or K=32). B cells in the data set (orange) start with approximately 15% of their measured genes overlapping with the “B.cell.naive signature”. After data smoothing the gene set overlap grows to over 30%.

### 2. Grouping cells with the “groupby”

When analyzing a combined or integrated dataset (e.g. contains several samples, patients, or batches), it may be beneficial to first group cells into phenotypically similar subsets before building the KNN graphs. The “groupby” parameter, is a list that maps to a set of categorical variables in the AnnData.obs table, and is used to sort the cells into chunks that can be processed in parallel. We use python’s multiprocessing starmap function to asynchronously process each “groupby-group”. This leads to a set of smoothed count matrices that are each specific to a selected phenotype. For example, one might group cells by cluster label, so smoothing is constrained to transcriptionally similar cells.

### 3. The nearest neighbor graph

After grouping, a nearest neighbor graph of cells is constructed using scanpy.pp.neighbors, which in turn uses PyNNDescent [17]. To calculate distances between cells and to determine nearest neighbors, we use a density-adjusted Gaussian kernel, commonly used in graph based clustering and dimension reduction of scRNAseq [18]. The choice of K in building the KNN, will be context dependent, but in practice, setting K to around 32 works well. One suggestion to determine K, is to use the ‘geneset overlap’ function, which returns the size of the intersection between the gene set and the expressed genes (above a given threshold). When the ‘geneset overlap’ score plateaus, the limitations of the data become apparent (see Figure 1).

### 4. Matrix smoothing

To address the noisy and sparse nature of single cell expression data, we apply nearest neighbor smoothing to produce a smoothed gene expression profile for each cell based on its neighbors. This assumes that gene expression varies smoothly along the data manifold (approximated by the nearest neighbor graph) and hence we can use information from neighboring cells to denoise the expression profiles of cells (similar to Gaussian smoothing in images, which assumes that pixel intensities vary smoothly in space) [19, 20]. The smoothed expression matrix of cells by genes is calculated via matrix multiplication *AX* = *M* where *A* represents a binary adjacency matrix or a weighted matrix of connectivities, and *X* is the cell by gene matrix of gene expression counts. Additionally, one can set the ‘smooth mode’ to ‘off’ in order to disable smoothing. As both A and X are typically sparse, our implementation uses the scipy sparse matrix library as an effort to be mindful of memory use [21].

### 5. Scoring Functions

As part of the package, several functions are available for gene set scoring. These include ‘singscore’ [6], ‘ssgsea’ [5, 22], ‘rank biased overlap’ [23], ‘mean z score’, ‘average score’, ‘median score’, ‘summed up’ [24] and a ‘geneset overlap’ count. For each gene set, the gene set scores are recorded in the AnnData.obs pandas table, with one column per gene set, facilitating visualization through compatibility with the Scanpy plotting system.

### 6. Validation of GSSNNG

In order to validate the method, we used three data sets with known ‘ground truths’. First, we compare smoothed and non-smoothed gene set scoring to identify B cells in a mixture of PBMCs[25]. Second, we show how smoothed gene set scoring helps to identify cells with an immune response in a dataset of phagocytes treated with LPS [8]. Third, we assess smoothed and non-smoothed gene set scoring of cellular response genes in a dataset of endothelial cells exposed to spinal injury in mice [26].

### 6.1. 10X genomics pbmc3k

In this data set, peripheral blood mononuclear cells (PBMCs) from a healthy donor were sequenced and annotated with cell type labels. The pre-processed data contains 2,638 cells with 1,838 genes. We applied the LM22 gene sets (cell type signatures) to produce cell type specific scores [27], focusing on the naive B cell signature (Figure 1B). The ‘geneset overlap’ function was applied to non-smoothed data and smoothed data (K=8 or K=32 neighbors). This function returns the number (or fraction) of genes that have expression measures above the given threshold. Here we show that for previously annotated B cells the number of genes from the B.cell.naive signature was effectively doubled with smoothing. With the limited set of genes available, the increase in gene set overlap does not improve noticeably past 32 neighbors. But clearly, through the use of matrix smoothing, the B cell specific signal was improved without an effect to non-B cells and allows one to easily distinguish B-cells from other cells using the signature scores.

### 6.2. Hagai et al

Murine phagocytic cells were treated with lipopolysaccharide (LPS) which causes a strong immune response. The OKUMURA INFLAMMATORY RESPONSE LPS gene set [28] contains genes related to this cellular warning system (see Figure 2). The data set was subsampled to 2,340 lps4 treated cells and 2,104 untreated cells and scored using the ‘summed up’ ranks function, which simply sums up the smoothed and ranked expression of signature genes for each cell. The results are compared between no smoothing and smoothing with 4, 8, or 32 neighbors. It was observed that with increasing neighborhood size, the separation in the distribution of gene set scores increased, improving the prediction of treatment group (Figure 2). Using sklearn’s roc auc score function, the area under the curve (AUC) was calculated to be 0.95 for unsmoothed data, and improved to 1.0 for smoothed data.

**Fig. 2.**
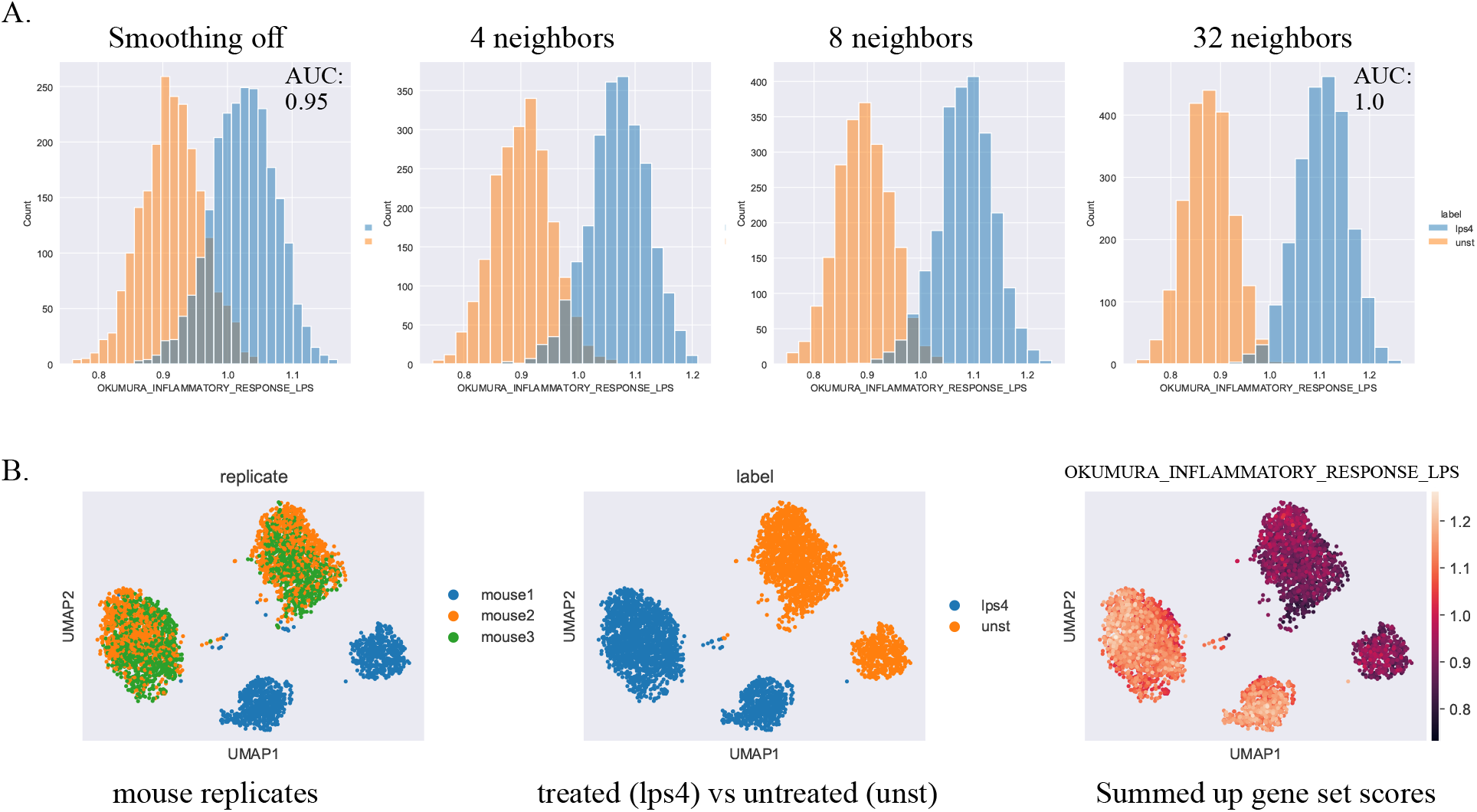
Figure 2. (A) Murine phagocytic cells were scored using the “Okumura inflammatory response LPS” gene set and the ‘summed up’ function. The histogram shows that by increasing the size of the neighborhood, the gene set score distributions separate between treated and untreated groups. The AUC was improved from 0.95 to 1.0 with smoothing. (B) UMAP plot of the cells in (A).

### 6.3. Squair et al

In this experiment, mice received spinal injuries which provoked a cellular response. The study reported that from all cells investigated, endothelial cells showed the greatest response. From the differential expression analysis on endothelial cells, 19 genes were validated using RNAscope. Because not all genes were validated, the gene sets were constructed from 12 of the 19 genes, with 7 showing higher expression in the injured mice and 5 showing lower expression for injured mice, thus making a two part gene set (both up and down). Gene set scoring was performed using the ‘rank biased overlap (RBO)’ and ‘ssGSEA’ functions on ranked expression on endothelial cells, and it was observed that with smoothing the gene set score distributions became separated, more clearly defining the the injured and the control groups (Figure 3). In predicting exposure, the area under the curve (AUC) was 0.62 and 0.69 using unsmoothed data, which improved to 0.71 and 0.8 with smoothed data after applying the RBO and ssGSEA score functions respectively.

**Fig. 3.**
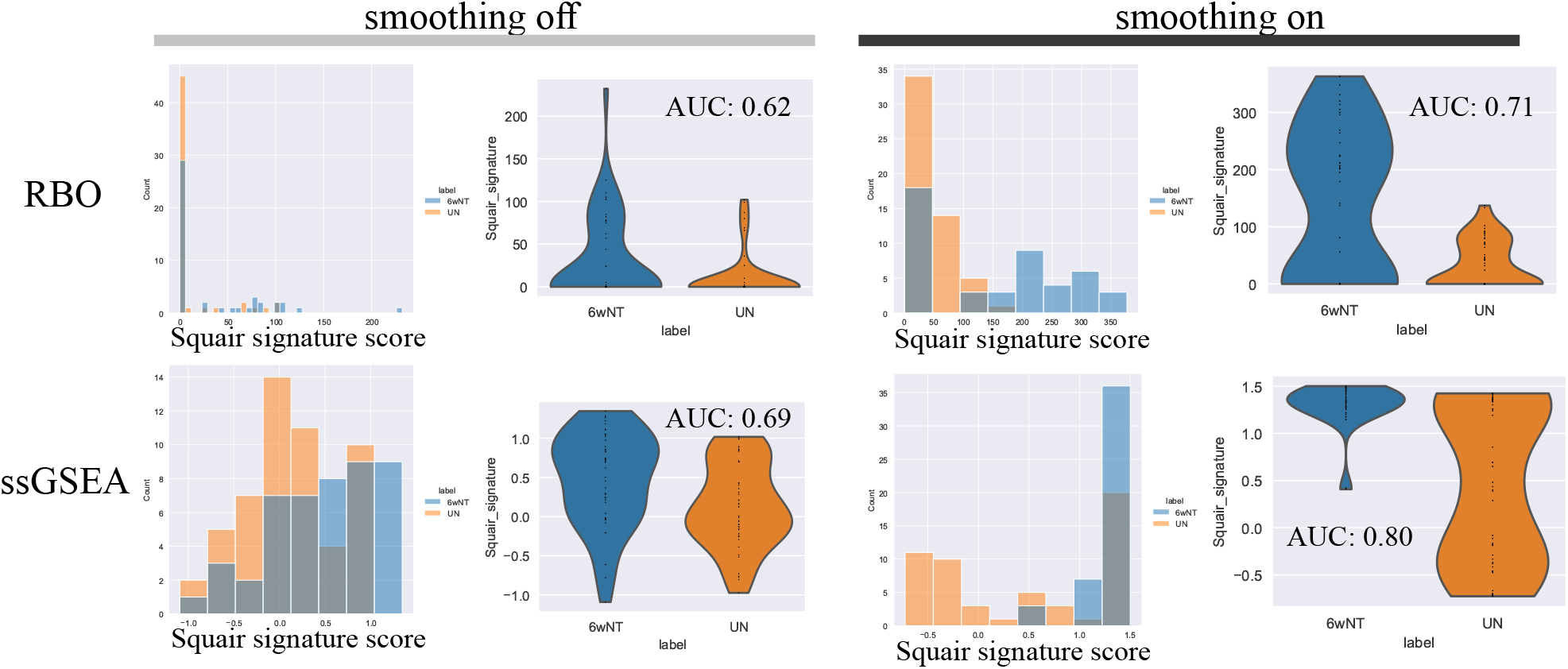
Figure 3. Endothelial cells from mice who received spinal injuries show a response to hypoxic environment. Scores are shown from no smoothing and smoothing with 32 neighbors using two methods, rank biased overlap (RBO) and single sample GSEA (ssGSEA). The AUC in predicting the cell label was improved with both scoring functions (0.62 to 0.71 for RBO and 0.69 to 0.80 for ssGSEA).

## 7. Availability

The gssnng package is available on github (https://github.com/IlyaLab/gssnng) and the PyPI index (https://pypi.org/project/gssnng/).

## Competing interests

No competing interest is declared.

## Author contributions statement

D.G. and M.S. conceived of, and implemented, the method.

D.G. and M.S. produced and analysed the results. D.G., M.S.,

S.H. wrote and reviewed the manuscript.

## Acknowledgments

This work was supported by a Cancer Research UK Grand Challenge (CRUK grant 29068). Additional support from the Tlsty Lab at UCSF (PI Tlsty, CRUK grant 27145), McGill University Thoracic and Upper GI Cancer Research Laboratories (PI Ferri, CRUK grant 29071) and the Advanced Genomic Technologies Laboratory (PI Ragoussis, CRUK grant 29078) is greatly appreciated.

## Notes

https://github.com/Gibbsdavidl/gssnng

### Competing Interest Statement

The authors have declared no competing interest.

### Summary of Updates

Text has been expanded with all new figures and new references.

## References

1. Aravind Subramanian, Pablo Tamayo, Vamsi K Mootha, Sayan Mukherjee, Benjamin L Ebert, Michael A Gillette, Amanda Paulovich, Scott L Pomeroy, Todd R Golub, Eric S Lander, et al. Gene set enrichment analysis: a knowledge-based approach for interpreting genome-wide expression profiles. Proceedings of the National Academy of Sciences, 102(43):15545–15550, 2005.

2. Jui-Hung Hung, Tun-Hsiang Yang, Zhenjun Hu, Zhiping Weng, and Charles DeLisi. Gene set enrichment analysis: performance evaluation and usage guidelines. Briefings in bioinformatics, 13(3):281–291, 2012. Publisher: Oxford University Press.

3. Henryk Maciejewski. Gene set analysis methods: statistical models and methodological differences. Briefings in Bioinformatics, 15(4):504–518, July 2014.

4. Farhad Maleki, Katie Ovens, Daniel J. Hogan, and Anthony J. Kusalik. Gene Set Analysis: Challenges, Opportunities, and Future Research. Frontiers in Genetics, 11, 2020.

5. David A. Barbie, Pablo Tamayo, Jesse S. Boehm, So Young Kim, Susan E. Moody, Ian F. Dunn, Anna C. Schinzel, Peter Sandy, Etienne Meylan, Claudia Scholl, Stefan Fröohling, Edmond M. Chan, Martin L. Sos, Kathrin Michel, Craig Mermel, Serena J. Silver, Barbara A. Weir, Jan H. Reiling, Qing Sheng, Piyush B. Gupta, Raymond C. Wadlow, Hanh Le, Sebastian Hoersch, Ben S. Wittner, Sridhar Ramaswamy, David M. Livingston, David M. Sabatini, Matthew Meyerson, Roman K. Thomas, Eric S. Lander, Jill P. Mesirov, David E. Root, D. Gary Gilliland, Tyler Jacks, and William C. Hahn. Systematic RNA interference reveals that oncogenic KRAS-driven cancers require TBK1. Nature, 462(7269):108–112, November 2009.

6. Momeneh Foroutan, Dharmesh D. Bhuva, Ruqian Lyu, Kristy Horan, Joseph Cursons, and Melissa J. Davis. Single sample scoring of molecular phenotypes. BMC bioinformatics, 19(1):404, November 2018.

7. Tae Hyun Kim, Xiang Zhou, and Mengjie Chen. Demystifying “drop-outs” in single-cell UMI data. Genome Biology, 21(1):196, August 2020.

8. Jordan W. Squair, Matthieu Gautier, Claudia Kathe, Mark A. Anderson, Nicholas D. James, Thomas H. Hutson, Rémi Hudelle, Taha Qaiser, Kaya J. E. Matson, Quentin Barraud, Ariel J. Levine, Gioele La Manno, Michael A. Skinnider, and Grégoire Courtine. Confronting false discoveries in single-cell differential expression. Nature Communications, 12(1):5692, September 2021. Number: 1 Publisher: Nature Publishing Group.

9. Yael Baran, Akhiad Bercovich, Arnau Sebe-Pedros, Yaniv Lubling, Amir Giladi, Elad Chomsky, Zohar Meir, Michael Hoichman, Aviezer Lifshitz, and Amos Tanay. Metacell: analysis of single-cell rna-seq data using k-nn graph partitions. Genome biology, 20(1):1–19, 2019.

10. Aaron T L Lun, Karsten Bach, and John C Marioni. Pooling across cells to normalize single-cell rna sequencing data with many zero counts. Genome biology, 17(1):1–14, 2016.

11. Florian Wagner, Yun Yan, and Itai Yanai. K-nearest neighbor smoothing for high-throughput single-cell rna-seq data. BioRxiv, page 217737, 2017.

12. David van Dijk, Roshan Sharma, Juozas Nainys, Kristina Yim, Pooja Kathail, Ambrose J. Carr, Cassandra Burdziak, Kevin R. Moon, Christine L. Chaffer, Diwakar Pattabiraman, Brian Bierie, Linas Mazutis, Guy Wolf, Smita Krishnaswamy, and Dana Pe’er. Recovering Gene Interactions from Single-Cell Data Using Data Diffusion. Cell, 174(3):716–729.e27, July 2018. Publisher: Elsevier.

13. Melania Franchini, Simona Pellecchia, Gaetano Viscido, and Gennaro Gambardella. Single-cell gene set enrichment analysis and transfer learning for functional annotation of scrna-seq data. NAR Genomics and Bioinformatics, 5(1):lqad024, 2023.

14. F Alexander Wolf, Philipp Angerer, and Fabian J Theis. Scanpy: large-scale single-cell gene expression data analysis. Genome biology, 19:1–5, 2018.

15. Pau Badia-i Mompel, Jesús Vélez Santiago, Jana Braunger, Celina Geiss, Daniel Dimitrov, Sophia Muüller-Dott, Petr Taus, Aurelien Dugourd, Christian H Holland, Ricardo O Ramirez Flores, et al. decoupler: ensemble of computational methods to infer biological activities from omics data. Bioinformatics Advances, 2(1):vbac016, 2022.

16. Dénes Tuürei, Tamás Korcsmáros, and Julio Saez-Rodriguez. Omnipath: guidelines and gateway for literature-curated signaling pathway resources. Nature methods, 13(12):966–967, 2016.

17. Wei Dong, Charikar Moses, and Kai Li. Efficient k-nearest neighbor graph construction for generic similarity measures. In Proceedings of the 20th International Conference on World Wide Web, WWW ‘11, page 577–586, New York, NY, USA, 2011. Association for Computing Machinery.

18. Leland McInnes, John Healy, and James Melville. Umap: Uniform manifold approximation and projection for dimension reduction. arXiv preprint arXiv:1802.03426, 2018.

19. Linda G Shapiro and George C Stockman. Computer vision, volume 3. Prentice Hall New Jersey, 2001.

20. Jonathan Ronen and Altuna Akalin. netSmooth: Network-smoothing based imputation for single cell RNA-seq. F1000Research, 7:8, July 2018.

21. Etienne Becht, Leland McInnes, John Healy, Charles-Antoine Dutertre, Immanuel WH Kwok, Lai Guan Ng, Florent Ginhoux, and Evan W Newell. Dimensionality reduction for visualizing single-cell data using UMAP. Nature biotechnology, 37(1):38–44, 2019. Publisher: Nature Publishing Group.

22. Mohamed E. Abazeed, Drew J. Adams, Kristen E. Hurov, Pablo Tamayo, Chad J. Creighton, Dmitriy Sonkin, Andrew O. Giacomelli, Charles Du, Daniel F. Fries, Kwok-Kin Wong, Jill P. Mesirov, Jay S. Loeffler, Stuart L. Schreiber, Peter S. Hammerman, and Matthew Meyerson. Integrative radiogenomic profiling of squamous cell lung cancer. Cancer Research, 73(20):6289–6298, October 2013.

23. William Webber, Alistair Moffat, and Justin Zobel. A similarity measure for indefinite rankings. ACM Transactions on Information Systems, 28(4):20:1–20:38, November 2010.

24. Frédéric Pont, Marie Tosolini, and Jean J Fournié. Single-Cell Signature Explorer for comprehensive visualization of single cell signatures across scRNA-seq datasets. Nucleic Acids Research, 47(21):e133, December 2019.

25. 3k PBMCs Single Cell Gene Expression Dataset by Cell Ranger 1.1.0 from 10x Genomics.

26. Tzachi Hagai, Xi Chen, Ricardo J Miragaia, Raghd Rostom, Tomás Gomes, Natalia Kunowska, Johan Henriksson, Jong-Eun Park, Valentina Proserpio, Giacomo Donati, et al. Gene expression variability across cells and species shapes innate immunity. Nature, 563(7730):197–202, 2018.

27. Binbin Chen, Michael S Khodadoust, Chih Long Liu, Aaron M Newman, and Ash A Alizadeh. Profiling tumor infiltrating immune cells with cibersort. Cancer Systems Biology: Methods and Protocols, pages 243–259, 2018.

28. Shigeru Okumura, Jun-ichi Kashiwakura, Hisashi Tomita, Kenji Matsumoto, Toshiharu Nakajima, Hirohisa Saito, and Yoshimichi Okayama. Identification of specific gene expression profiles in human mast cells mediated by toll-like receptor 4 and fceri. Blood, 102(7):2547–2554, 2003.

